# Eco-Microbiology: Discovering Biochemical Enhancers of PET Biodegradation by *Piscinibacter sakaiensis*

**DOI:** 10.1101/2024.07.01.601593

**Authors:** Felipe-Andrés Piedra, Miguel A. Salazar, Raayed Rahman, Justin C. Clark, Anthony Maresso

## Abstract

The scale of plastic pollution boggles the mind. Nearly 400 megatons of virgin plastics are produced annually, with an environmental release rate of 80 percent; and plastic waste including micro- and nanoplastics are associated with a plethora of problems. The naturally evolved abilities of plastic-degrading and consuming microbes offer a starting point for generating sustainable and eco-centric solutions to plastic pollution. Here we developed an iterative discovery procedure coupling faster quasi-high-throughput polyethylene terephthalate (PET) dependent bioactivity screens with longer-term PET biodegradation assays to find small molecule and ionic boosters of PET consumption by the bacterium *Piscinibacter sakaiensis*. We discovered multiple hits supporting greater than 2-fold enhancement of PET biodegradation – with hits belonging to a small but heterogeneous set of compounds and mixtures, suggesting upregulation of PET consumption via multiple paths. This work has the potential to advance the creation of a fermentation-based process for solving PET plastic pollution.

**Importance:** Plastic pollution is an urgent environmental issue. In addition, micro- and nanoplastics (MNPs) have become an acute source of worry with discoveries of the global distribution and transport of MNPs, their presence within a diversity of organisms including common foodstuffs and human tissues, and their potential association with declining fertility and various disease states. Solutions are needed and the microbial world offers abundant help via naturally evolved biodegraders of plastic waste. We created a non-genetic method to accelerate polyethylene terephthalate (PET) plastic biodegradation by *Piscinibacter sakaiensis*, a bacterium that evolved to slowly but completely consume PET. Our method entails a combination of plastic-dependent bioactivity screens and slower biodegradation tests to find extrinsic biochemical stimulators of PET biodegradation. The conditions we found boost PET biodegradation by over two-fold and provide a foundation for further studies to realize a fermentation-based process needed to solve PET plastic pollution.

## Introduction

Plastics remain a revolutionary material, offering an extraordinary range and combination of desirable properties. However, their durability combined with a massive and accelerating scale of production means that plastic pollution has become a global and rapidly growing problem^1–3^.

In their roughly seven decades of existence, over eight gigatons of plastic have been released into the global environment; and the current annual rate of release is approximately 0.3 gigatons and growing^1^. Bulk plastic pollution creates a series of problems: as a sign of community and environmental neglect, and also by virtue of its association with runaway climate change, it contributes to a widespread feeling of dismay^4^; it can cause a plethora of infrastructure issues, especially in water management and flood control^5,6^; and it can be straightforwardly problematic to wildlife, causing injury or death to large organisms, especially marine life, via physical capture or accidental ingestion. In addition, bulk plastic pollution weathers and degrades into smaller debris and micro- and nanoplastics (MNPs); and it is now well established that MNPs, like other forms of small particulate matter, circulate globally through a variety of geophysical processes including atmospheric transport through the free troposphere^7,8^. MNPs also readily enter a staggering variety of organisms and tissues, including some of the most common fruits and vegetables consumed by human beings and both male and female human reproductive organs^7,9–13^. And once within an organism, a variety of pathologies become possible, whether due to the materials themselves (ex: contributing to the blockage of small arteries) or by virtue of the ability of MNPs to adsorb and deliver other chemicals including a plethora of plastic additives of varying toxicity^3,7,11,13–17^.

Addressing plastic pollution is therefore of vital concern, and it is also not surprising that a variety of microbes have already started^18,19^. Indeed, all plastics are polymeric hydrocarbons that can serve as a carbon source for microbes able to degrade them, notwithstanding other well known (ex: substrate for biofilm formation) and potential microbial uses. The major issue remains the scale and speed of the generation of plastic pollution, which demands both the adoption of sustainable patterns of production and consumption and cleaning up at a planetary scale. For the latter, the naturally evolved abilities of plastic-degrading microbes can be tapped to create low-cost technologies for the rapid and widespread biodegradation of plastic waste.

One such microbe is the recently described aerobic gram-negative bacterium, *Piscinibacter sakaiensis* (*P. sakaiensis)*^19^. *P. sakaiensis* evolved to degrade and completely consume the plastic polyethylene terephthalate (PET), which is a semi-aromatic polyester abundantly used in textiles, clothing, and food and beverage packaging. *P. sakaiensis* degrades PET into soluble fragments and further into its basic chemical building blocks of ethylene glycol and terephthalic acid, which are further catabolized and completely incorporated into the organism itself, either as basic biochemical constituents (nucleic and amino acids, lipids, etc.) or into another polymeric form, polyhydroxyalkanoate (PHA), which is presumably used as a carbon/energy source by *P. sakaiensis* when needed^19–21^. PHAs are readily degraded and consumed by other microbes^22^. The first and most important step in the biodegradation of PET by *P. sakaiensis* is accomplished by a single secreted enzyme, the IsPETase, which cleaves the ester bonds of the polymer into BHET and MHET and acts further on BHET to produce ethylene glycol and terephthalic acid (MHET is further degraded by a surface membrane bound MHETase) ^19^. The IsPETase has been intensely studied and its activity and stability considerably improved via a combination of rational and agnostic mutagenesis strategies^23,24^. However, *P. sakaiensis* has been largely ignored and little to no effort has been made to improve its extraordinary but quantitatively modest ability (∼0.1 mg·cm^2^·day^-1^) to completely consume PET, yielding biomass, CO_2_, and water as byproducts.

We developed a quasi-high-throughput assay to measure PET-dependent microbial bioactivity under hundreds of distinct chemical conditions, hypothesizing that a boost would lead to enhanced PET consumption by *P. sakaiensis*. Results from our downstream PET biodegradation tests generally supported our hypothesis, but also revealed surprising variation, especially in week-to-week biodegradation rates, establishing the need to generate an iterative process combining more rapid screening with longer PET biodegradation assays. Ultimately, we discovered several hits, especially in tandem, that enhanced total PET biodegradation by a factor of two or more. High dilutions of a rich medium alone and in combination with single chemical species belonging to a small but diverse set of compounds produced robust enhancement of PET biodegradation. In addition, a combination of sodium phosphate and ethylene glycol strongly enhanced total PET biodegradation while stimulating increasing rates of biodegradation. The heterogeneous nature of the enhancing compounds suggests the existence of multiple paths to upregulating PET consumption. Crucially, the conditions discovered are inexpensive and easily obtained. Thus, our work provides a foundation for further efforts to generate a fermentation-based process for the rapid conversion of PET waste to biomass using a small-molecule-stimulation effect as the prime driver of enhanced degradation.

## Results

### Phenotype microarray screening for enhanced PET-dependent bioactivity

Bacteria are highly adaptable, combining the constant sensing of environmental stimuli including discrete chemical cues with manifold metabolic and gene regulatory responses. This adaptability can be exploited experimentally, and tuned towards a desired behavior. Thus, we hypothesized the existence and discoverability of small molecules and ionic compounds able to upregulate the naturally-evolved propensity of *P. sakaiensis* to consume polyethylene terephthalate (PET).

Further hypothesizing that *P. sakaiensis* can be readily screened for novel activators of PET biodegradation, we used Biolog phenotype microarray (PM) plates to search chemical space for conditions stimulating PET-dependent bioactivity, with bacterial growth and metabolism as readouts (Fig. 1A). PET-dependent bacterial growth can report directly on the conversion of PET to microbial biomass; and PET-dependent metabolic activity, measured by adding a redox dye to the PM microcultures and recording the resultant color depth after a 24 hr incubation, should report on the level of work performed by *P. sakaiensis*, a quantity that may be associated with elevated biodegradation. Each aspect of PET-dependent bioactivity was quantified using a *synergy score*, with values in excess of one indicating a non-linear increase in bacterial growth or metabolism from the relevant chemical:PET combination (Fig. 1B)(see Methods).

**Figure 1.**
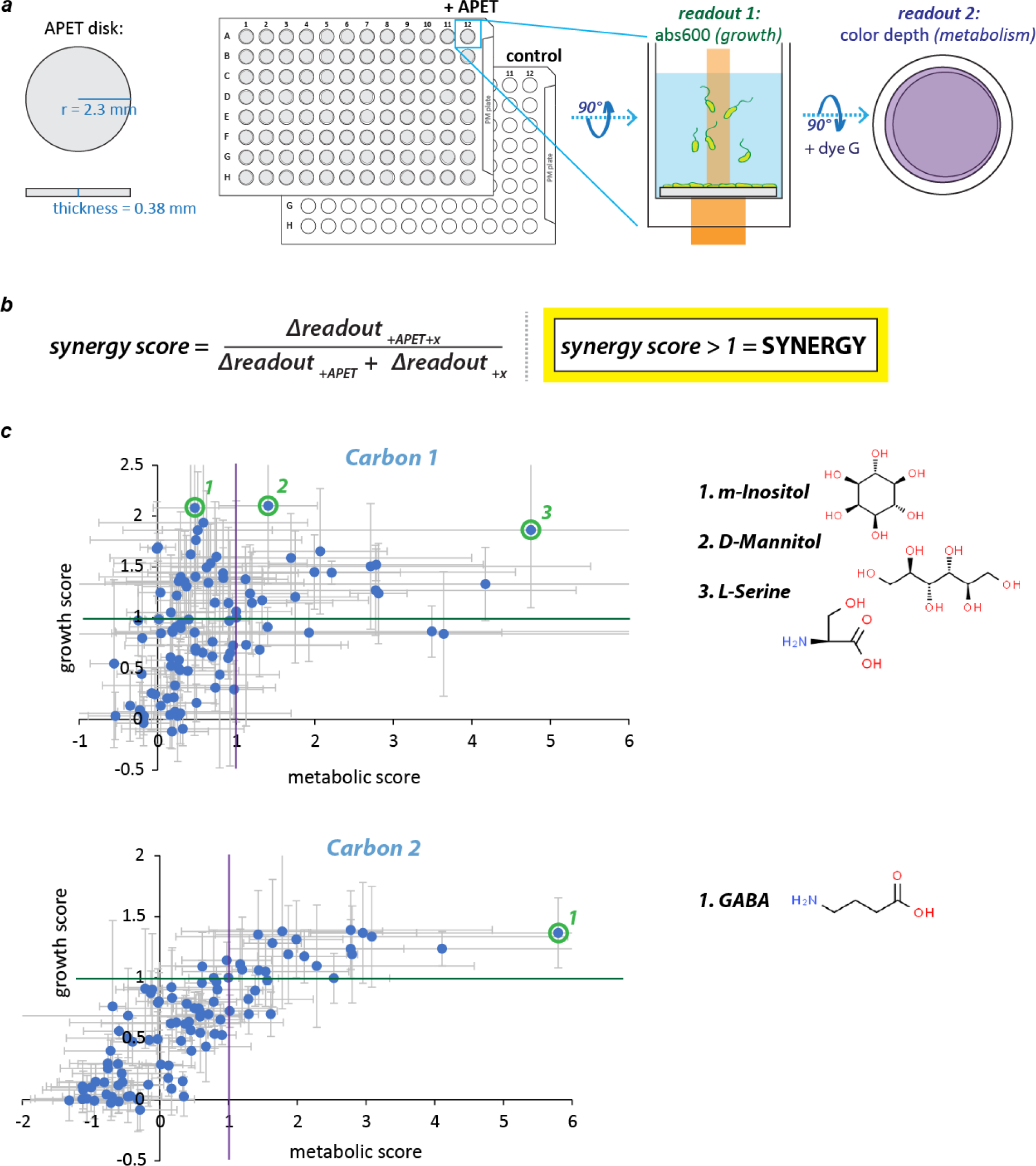
Screening for *synergy* or enhancement of PET-dependent growth and metabolism in phenotype microarrays. **(A)** A microplate-based assay for finding chemical conditions supporting enhanced PET-dependent growth and metabolism of *P. sakaiensis*. Phenotype microarrays (PM plates; Biolog) were prepared with and without disks of amorphous polyethylene terephthalate (PET), loaded with *P. sakaiensis* in YSV minimal medium, and incubated at 30°C with shaking for 4 days. Bacterial growth measurements were made every hour or once at the end of the experiment by recording A600s from each well. Metabolic activity was measured at a single time-point following the introduction of a redox dye (dye G) on day 4 and a 24 hr incubation. **(B)** Synergy scores. The two readouts were used to calculate synergy scores reflecting the presence (>1) or absence (<=1) of enhanced PET-dependent growth and/or metabolic activity (see Methods). Enhancements of either, but especially growth, were hypothesized to promote enhanced PET biodegradation. **(C)** Carbon source plates (PM1 and PM2A) showed correlated growth and metabolic scores, with a minority of conditions supporting both growth and metabolic synergy. Each plot contains 96 data points (95 test conditions + 1 no chemical control per PM plate) where each data point represents the mean of 3 independent experiments (error bars = standard dev). Carbon 1 = PM1 plate; Carbon 2 = PM2A plate.

The full spectrum of possible bioactivity levels can be decomposed into four major quadrants (Fig. 1C and 2): 1) upper right conditions (g and m scores greater than one) support enhanced PET-dependent bacterial growth and metabolism; 2) upper left conditions (only g scores greater than one) support greater than expected growth but less than expected metabolic activity; 3) lower left conditions (g and m scores less than one) support diminished growth and metabolic activity; and 4) lower right conditions (only g scores less than one) support diminished growth with greater than expected metabolic activity. Only the upper quadrants are potentially supportive of enhanced PET consumption, so we focused on ‘hits’ from within these two regions. Moreover, we prioritized hits available for ≤ $1 per gram for further study.

**Figure 2.**
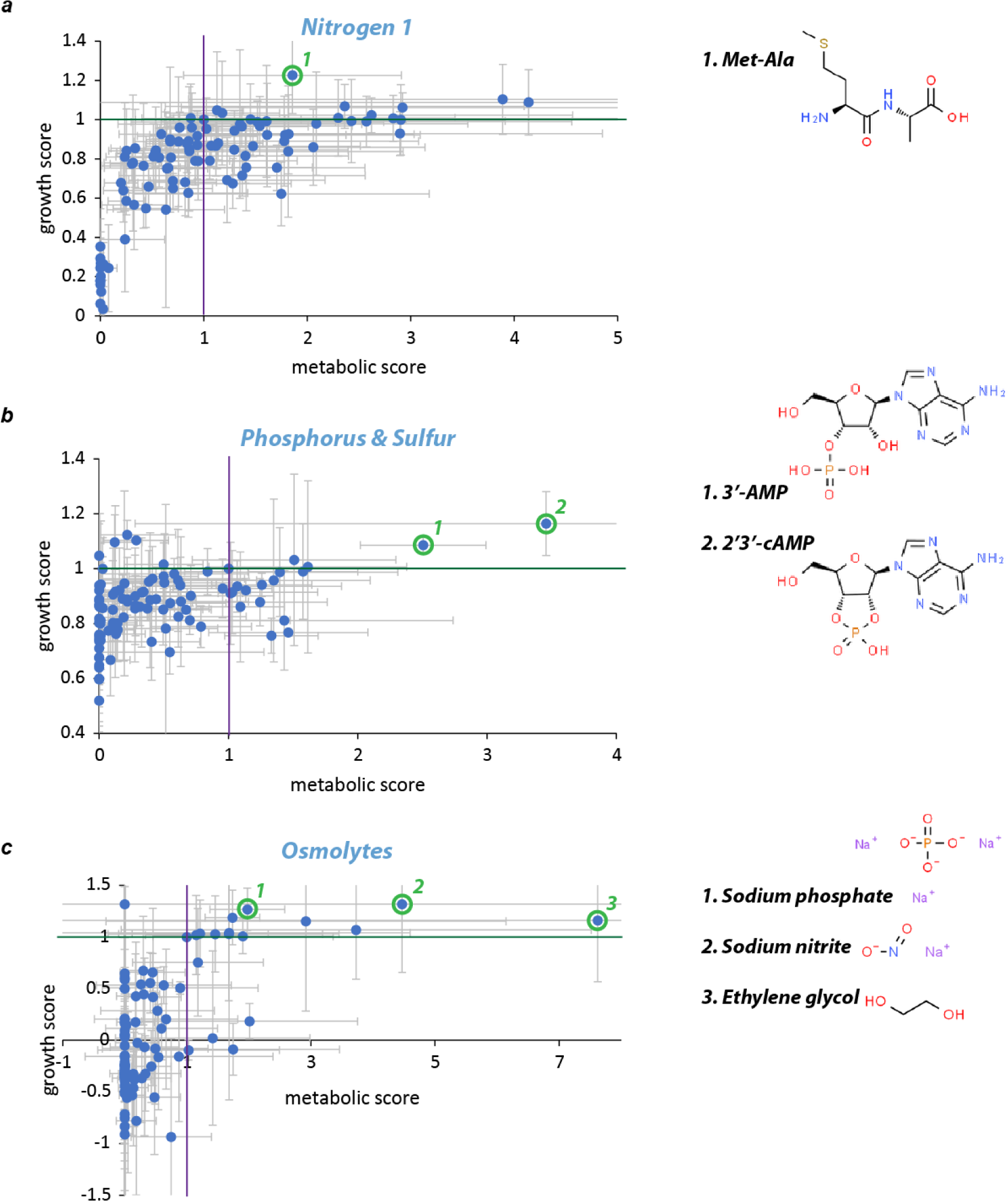
Few nutrient sources and inorganic compounds support both growth and metabolic synergies. **(A)** Nitrogen sources (PM3B) showed little synergy, with a dipeptide (Met-Ala) modestly boosting PET-dependent growth and metabolism. **(B)** Phosphorus and sulfur sources (PM4A): two nucleotide derivatives supported growth and metabolic synergy. **(C)** Osmolytes (PM9): two inorganic compounds and one byproduct of *P. sakaiensis* IsPETase activity supported growth and metabolic synergy. Each plot contains 96 data points (95 test conditions + 1 no chemical control) where each data point represents the mean of 3 independent experiments (error bars = standard dev).

### Searching for hit concentrations supporting enhanced PET-dependent growth

Chemical quantities in PM plates were not given, so it was necessary to determine the concentrations of hits supporting enhanced PET-dependent bioactivity. In addition, anticipating the need to scale up from 150 microL cultures to larger volumes needed for weeks-long PET biodegradation experiments, we assayed bacterial growth scores across a 32-fold range of hit concentrations in a larger 24-well format (1.05 ml per well); absorbance measurements were made after two days instead of four because of enhanced evaporative loss from greater air-exposed surface area and fluid mixing in the larger wells. Finally, we included dilutions of a rich culture medium (growth medium #802, or gm802) as an eighth possible hit. Using these conditions and scoring growth as described above, only seven of 48 conditions assayed supported synergy scores greater than one (Fig. 3A). The six highest growth scores (range: 1.68 to 2.22) all came from dilutions of gm802, with the seventh highest coming from the lowest percentage (0.016%) of L-serine assayed (Fig 3A).

**Figure 3.**
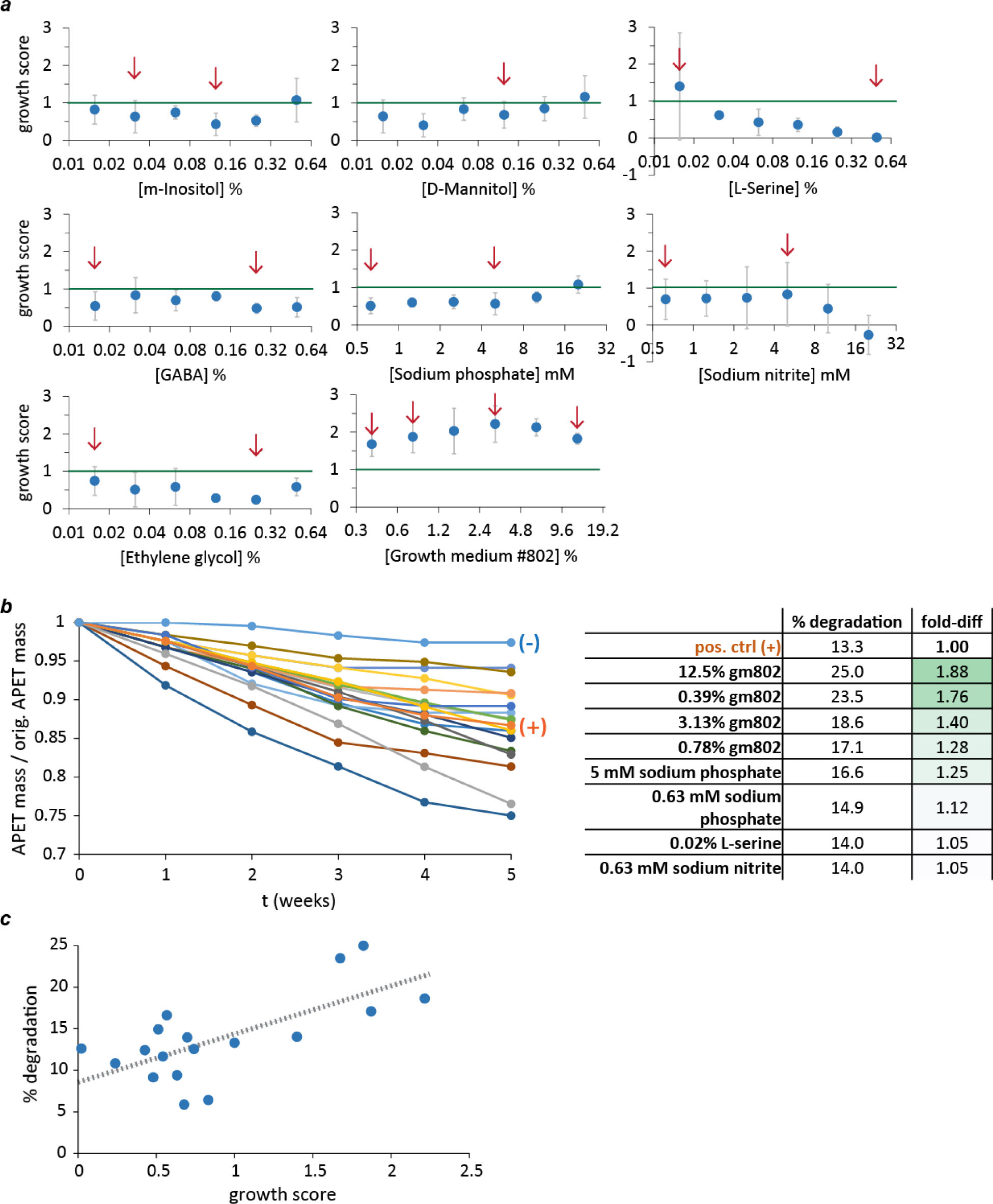
Higher growth scores correlate with faster PET biodegradation. **(A)** Growth scores over a range of concentrations of PM screen-identified hits and dilutions of growth medium #802 (gm802). Chemical conditions were prepared across a 32-fold range of concentrations and assayed in 24-well plates for growth scores after 2 days. Each data point represents the mean of 3 independent experiments (error bars = standard dev). Data points marked with a red arrow indicate conditions chosen for a downstream PET biodegradation assay. **(B)** Dilutions of growth medium #802 and 3 other chemicals improved PET biodegradation. A 5-week-long PET biodegradation assay (see Methods) was performed using 17 test conditions and 2 unsupplemented YSV control conditions [(+) = with *P. sakaiensis*; (-) = no bacteria]. PET strip masses were measured weekly. Fold-differences in % biodegradation after 5 weeks (fold-diff) were calculated relative to the unsupplemented positive control **(C)** The extent of PET biodegradation after 5 weeks correlated with the growth scores measured after 2 days. (Linear fit: y = 5.48x + 8.80; R^2^ = 0.45).

### Assessing single hits for enhanced PET biodegradation

We chose a subset of 17 conditions including high and low growth score extremes to scale up for PET biodegradation assays needed to determine the effectiveness of our bioactivity-based screening. Larger cultures (4 ml) were used to accommodate plastic strips large enough for accurate weekly measurements of PET mass loss. Each culture contained two strips of PET plastic differing only slightly in shape (straight vs. round edges). Every week, one strip was removed and processed (1% SDS wash, 70% ethanol wash, drying) before being weighed, and 90-95% of the media was replaced with fresh media. The purpose of this strategy was to minimize disruption of biodegradation by ensuring that each culture could always contain a population of *P. sakaiensis* in a PET-consuming biofilm. Biodegradation rates (mg·cm^-2^·day^-1^) were measured from the weekly change in mass and used to estimate total mass loss in each of the conditions assayed (Fig. 3B).

Remarkably, the extent of biodegradation measured at five weeks correlated with the average growth scores measured after two days (Fig. 3C). An average PET biodegradation rate of 0.096 ±0.037 mg·cm^2^·day^-1^ was measured for the positive control (YSV only) (Supp. Table 1), which is consistent with that reported by Yoshida et al^19^. A total of eight conditions supported equal or greater biodegradation than control, with the greatest gains coming from the four dilutions of gm802 tested (Fig 3B). The two conditions showing the greatest enhancement were the high and low concentration extremes of gm802 (+88% and +76% from 12.5% and 0.39% gm802, respectively); but the biodegradation rate from 12.5% gm802 slowed markedly between weeks four and five, while that of 0.39% showed no such slowing (Fig. 3B and Supp. Table 1). Conditions showing modestly enhanced biodegradation relative to control were 5 and 0.625 mM sodium phosphate pH 7 and 0.016% L-serine.

### Combining hits for greater enhancement of bacterial growth and PET biodegradation

We hypothesized that greater gains in PET-dependent bacterial growth and PET biodegradation could be realized by combining chemical conditions (Fig. 4). To that end, we started by searching a range of concentrations within 14 different pairwise combinations for novel conditions supporting growth scores greater than one (Fig. 4A). The 14 combinations came from two different starting concentrations of gm802 (12.5% and 0.39%) x four chemicals (= sodium phosphate, L-serine, GABA, and ethylene glycol) plus all six pairwise combinations generated from the four non-gm802 conditions. The dilution scheme was chosen to explore concentrations equal to, 2-fold higher, and 2-fold lower than those supporting or nearly supporting growth synergy for the single chemicals involved (Fig. 4A). Single chemical controls were also included in the screen (Fig. 4A).

**Figure 4.**
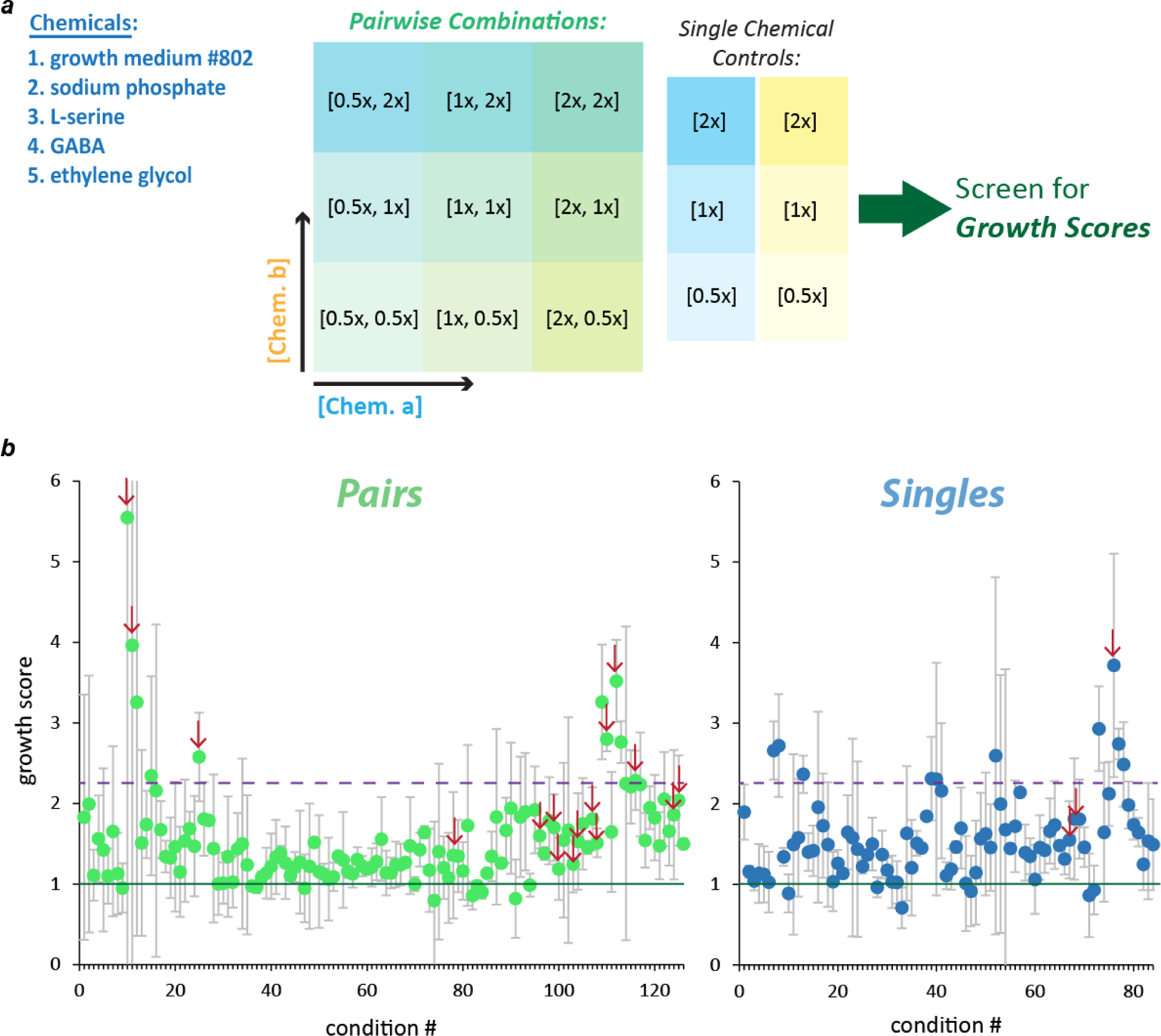
Screening pairwise combinations of choice chemicals for greater growth synergy. **(A)** Subset of hits chosen for screen of pairwise combinations. 5 different chemical conditions were chosen for 14 different pairwise combinations: 2 different starting concentrations of gm802 (12.5% and 0.39%) x 4 chemicals (= sodium phosphate, L-serine, GABA, and ethylene glycol) plus all 6 pairwise combinations generated from the 4 non-gm802 conditions. The dilution scheme was chosen to explore concentrations equal to, 2-fold higher, and 2-fold lower than those supporting or nearly supporting growth synergy for the single chemicals involved. Single chemical controls were also included. **(B)** Growth scores across pairwise combinations and single chemical controls. The vast majority of conditions tested supported growth synergy (= data points above green line) and 10 pairwise combinations + 9 single chemical controls showed growth synergy exceeding the maximum observed in the first set of single chemical 24-well-plate assays performed ( = data points above dashed purple line). Each data point represents the mean of 2 independent experiments (error bars = standard dev). Data points marked with a red arrow indicate conditions chosen for a downstream PET biodegradation assay.

Almost all growth scores were above one (92%), and 10 of the 126 pairwise combinations supported average growth scores higher than the max observed in the screen of single hits (Fig. 4B). With the exception of exceptionally high growth scores from combinations of sodium phosphate and ethylene glycol, combinations involving dilutions of gm802 generally supported the highest growth scores. We selected a subset of 19 test conditions including three of the highest growth scores and three gm802 controls for another 5-week-long PET biodegradation assay (Fig. 5). The average PET biodegradation rate in unsupplemented YSV medium was 0.043 ±0.011 mg·cm^2^·day^-1^, approximately half that of the first experiment (Supp. Table 2). Also contrary to the single hits data, almost all conditions showed enhanced biodegradation after 5 weeks (Fig. 5 and Supp. Table 2). A total of 10 conditions showed greater fold-enhancement than the previous max of 1.88, with six conditions supporting greater than 2-fold enhancement (Fig 5). Most of the 17 conditions that supported enhanced biodegradation were dilutions of gm802 in combination with either sodium phosphate, ethylene glycol, GABA, or L-serine, but three contained no rich medium at all: these were sodium phosphate with either ethylene glycol (2 conditions) or GABA (Fig 5). Consistent with experiment one, the biodegradation rate from 12.5% gm802 decreased significantly over the course of the experiment (Supp. Tables 1 and 2). In contrast, biodegradation rates from both combinations of sodium phosphate and ethylene glycol increased significantly over the five weeks assayed (Supp. Table 2).

**Figure 5.**
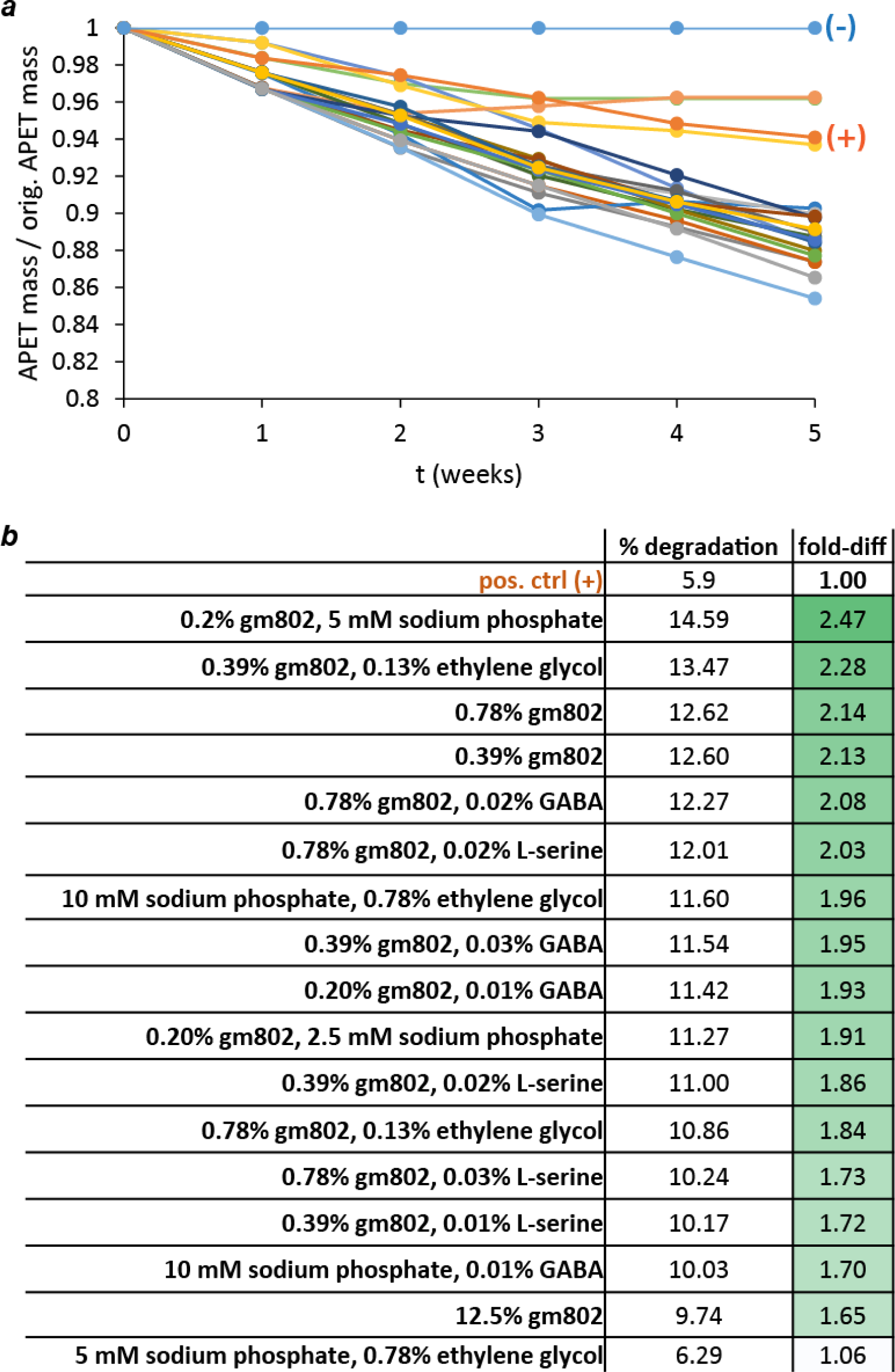
Pairwise combos support enhanced PET biodegradation. **(A)** A 5-week-long PET biodegradation assay (see Methods) was performed using 19 test conditions and 2 unsupplemented YSV control conditions [(+) = with *P. sakaiensis*; (-) = no bacteria]. Most conditions resulted in enhanced PET biodegradation. **(B)** The 17 test conditions supporting enhanced PET biodegradation. Fold-differences in % biodegradation after 5 weeks (fold-diff) were calculated relative to the unsupplemented positive control, with 6 conditions yielding 2-fold or higher enhancement.

## Discussion

We report a novel strategy based on extrinsic chemical stimulation to accelerate the consumption of PET plastic by a recently described and naturally evolved PET-eater, *Piscinibacter sakaiensis*. By creating a quasi-high-throughput PET-dependent bioactivity assay to screen a library of 475 chemical conditions and setting up an iterative discovery procedure combining more rapid screening with longer-term biodegradation assays, we discovered conditions supporting a greater than two-fold gain in PET biodegradation over that achieved by *P. sakaiensis* in YSV minimal medium alone. Crucially, the conditions themselves are simple and inexpensive, and therefore easily obtained in a variety of settings.

Multiple single hit and pairwise hit combinations were discovered that enhance both PET-dependent bacterial growth and PET biodegradation. Among these, low concentrations (<1%) of growth medium #802 (gm802), a rich medium similar to Luria-Bertani broth (LB), robustly enhanced both, and did so both alone and in combination with single members of a small but diverse set of chemical species (sodium phosphate, ethylene glycol, GABA, L-serine). Indeed, combinations of gm802 with sodium phosphate and, separately, ethylene glycol, showed the highest increases in total PET biodegradation. Finally, a combination of sodium phosphate and ethylene glycol without any gm802 also proved highly promising, supporting enhanced PET-dependent bacterial growth and increasing rates of PET biodegradation over the five weeks assayed.

Of the five Biolog phenotype microarray (PM) types used in our initial screening – two carbon source plates (PM1 and PM2A), one phosphorus and sulfur source plate (PM4A), one nitrogen source plate (PM3B), and one osmotic conditions plate (PM9) – the two carbon source plates contained the greatest number of conditions supporting enhanced PET-dependent bioactivity. These were mostly simple sugars and amino acids, substrates readily used and funneled through central carbon metabolism across domains of life, that are clearly in short supply in minimal medium. The number and diversity of these sugars and amino acids helps make clear how high dilutions of gm802, which contain an enormous diversity of similar biomolecules all at very low concentration, would greatly support enhanced PET-dependent bioactivity.

After our initial screening, we switched from the 96-well format of PM plates to 24-well plates to determine the concentrations of eight chosen hits (including dilutions of gm802) supporting enhanced PET-dependent bacterial growth. The scaling up was done to better mirror the conditions expected in the larger 4 ml cultures of our PET biodegradation assays (higher volumes were needed to accommodate and completely submerge the PET strips used for accurate weekly measurements of mass loss). An expected effect of moving from the 150 microL microcultures of PM plates to the 1.05 ml cultures of 24-well plates is increased aeration from the emergence of surface wave action when plates are shaken at the rpms used here (260-280). Given that *P. sakaiensis* is an aerobic microbe, this increased aeration might be expected to increase PET-dependent growth; however, to our mild surprise, most hits across the 32-fold range of concentrations tested failed to support bacterial growth synergy. It is possible that in PM plates the hits are at concentrations outside of our downstream screening. There might also be a boost in PET-dependent growth from increased aeration that exceeds, in most cases, the effect of the added compound. Moreover, all of this indicates the importance of optimizing the dissolved oxygen content in future experiments to accelerate PET consumption by *P. sakaiensis*.

Remarkably, average growth scores measured after two days in 24-well plates correlated with the extent of PET biodegradation measured after five weeks, indicating the soundness of our screening procedures (Supp. Fig 1). However, growth scores (and especially metabolic scores from our initial screening) show high variation. This variation likely came from multiple sources, including 1) low-level variation in the percent crystallinity of amorphous PET plastic used; 2) variation in bacterial state at the start of each experiment; 3) the effects of well position-dependent evaporative loss; and 4) random artifacts associated with measurement. It is hard to speculate on the magnitude of each, but future efforts will be made to minimize experimental noise.

The PET biodegradation assays containing single hits exclusively and pairwise combinations of hits showed both interesting overlap and important differences (Fig. 3 and 5, Supp. Tables 1 and 2). Average PET biodegradation rates from unsupplemented control (YSV only) and three dilutions of gm802 (0.39, 0.78, and 12.5%) decreased roughly 2-fold between the two assays, suggesting the occurrence of a global change (Supp. Tables 1 and 2). Indeed, we suspect an increased percent crystallinity of PET to be the cause. It is known that PET becomes more refractory to IsPETase-mediated biodegradation with increasing percent crystallinity^19,24,25^. We treated our PET plastic with 10% bleach (0.6% sodium hypochlorite) between the two assays; unknown to us then, sodium hypochlorite alters the surface of PET^26^. This treatment was performed to deal with a contaminant growing in the 70% ethanol within which the PET was stored. Future instances of this problem can be averted by storing PET plastic under dry sterile conditions.

Although the four controls in the two PET biodegradation assays differed systematically in average biodegradation rates, the rates show similar dynamics through time (Supp. Tables 1 and 2). Indeed, biodegradation rates from 12.5% gm802 dropped significantly over the course of both assays, and even more sharply in assay two; while the others showed much less change (Supp. Tables 1 and 2). A similar drop in the biodegradation rate was observed for 25% gm802 in another assay, but that condition did not support enhanced PET biodegradation (Supp. Table 3). This suggests that *P. sakaiensis* adapts out of the PET-biodegradation-boosting effects of low concentrations of rich medium over a certain range (> 0.78% and ≤ 12.5%) well before complete PET degradation, perhaps forming a non-degrading but growing biofilm maintained by flows of dissolved nutrients from the rich medium. This highlights the need to assess biodegradation beyond five weeks, and ideally to the point of maximal or total degradation.

Very low concentrations of gm802 (0.39% and 0.78%) and their combination with low concentrations of sodium phosphate, ethylene glycol, GABA, or L-serine all showed steady 2-fold or greater enhancement of PET biodegradation over five weeks. In a similar biodegradation assay performed before PET treatment with 0.6% sodium hypochlorite, 0.39% gm802 with a low concentration of L-serine (0.016%) showed over 4-fold enhancement after 5 weeks and resulted in close to 50% biodegradation (Supp. Table 3). This condition supported a comparatively modest 1.86-fold enhancement in biodegradation assay two (Fig. 5 and Supp. Table 2). This suggests that many of the conditions discovered in assay two are capable of greater than 4-fold enhancement of PET biodegradation absent PET pretreatment with sodium hypochlorite.

One combination of chemicals without gm802 showed remarkable promise as a *P. sakaiensis*-mediated PET consumption enhancer: sodium phosphate and ethylene glycol (Fig. 5 and Supp. Table 2). The two conditions tested supported enhanced biodegradation after five weeks and at rates that increased over the course of the assay, especially between weeks one and two (Supp. Table 2). The presence of sodium phosphate here and in biodegradation-enhancing combinations with gm802 suggests the importance of phosphate availability for *P. sakaiensis* while it degrades and consumes PET. The presence of ethylene glycol seems even more interesting because it is a byproduct of IsPETase action. This might suggest the existence of a cellular mechanism(s) coupling, in a free phosphate concentration-dependent way, the sensing of ethylene glycol with the expression of IsPETase. Future experiments exploring the effects of these conditions on *P. sakaiensis* gene expression are needed.

In total, our work represents an important step toward creating a fermentation-based process for the rapid conversion of PET waste to microbial biomass. The iterative screening methodology developed can also be adapted to the discovery of ideal conditions for microbes able to consume other forms of plastic pollution.

## Methods

### Piscinibacter sakaiensis cultivation

Lyophilized *Piscinibacter sakaiensis (P. sakaiensis)* was received from the Biological Resource Center (NBRC), Japan. The lyophilized bacteria were reconstituted according to NBRC’s instructions. An initial glycerol stock (25% glycerol) was made from freshly reconstituted *P. sakaiensis* and used to streak plates of 1.5% agar in growth medium #802 (gm802). Plates were incubated at 30°C for 4-5 days before the appearance of single colonies.

A second glycerol stock of larger volume was made for all experiments described here in the following manner: a single colony was used to inoculate 4 ml of YSV containing a strip of PET and another single colony was used to inoculate 4 ml of only YSV; these cultures were grown for 6 days at 30 °C with shaking (260-280 rpm) and checked for PET-dependent growth by measuring and comparing their optical densities (OD600); after confirming PET-dependent growth, 1 ml was taken from the PET-containing culture and used to inoculate 25 ml of gm802 within a 125 ml Erlenmeyer flask which was then grown at 30°C with shaking (280 rpm); the gm802 culture was grown to an OD600 of 1 and glycerol stocked by mixing the culture 1:1 with 50% glycerol in water and storing at -80°C in 1 ml aliquots.

For all experiments described here, *P. sakaiensis* ‘overnight’ cultures were prepared by inoculating thawed glycerol stock (treated as a 30x stock) directly into gm802 (3-15 ml cultures) and incubating at 30°C with shaking (260-280 rpm) for approximately 24 hrs. After growing, an OD600 was measured and an appropriate volume of culture was pelleted (1080xg for 10 minutes) and washed in YSV medium (2x) for use in screening or PET biodegradation assays. We settled on this procedure after observing considerable variation in the growth time needed for ‘overnight’ cultures inoculated with single colonies from plates (1.5% agar in gm802). Whether fresh colony-containing plates were stored at 4°C or RT, single colonies lost the ability to support planktonic growth in rich media after 2-3 weeks.

### Media preparation

YSV medium and growth medium #802 were prepared according to Yoshida et al.^19^

### Preparation, storage, and use of amorphous polyethylene terephthalate (PET)

Sheets of amorphous PET (11”x12”x0.015”) were ordered from Polymer Firms (Tyngsboro, MA, USA) and cut into disks or strips of variable size using a Cricut Maker craft cutter. PET pieces were stored in 70% ethanol and at 4°C. Before each experiment, PET pieces were removed from 70% ethanol and dried on plastic weigh dishes inside of a biosafety cabinet. PET pieces were then UV irradiated for 10 minutes. After UV irradiation, PET pieces were introduced one-by-one into Biolog PM plates, 24-well plates, or 15 ml conicals used in PET biodegradation assays.

### Biolog phenotype microarray (PM) and 24-well plate screening

For a single experiment, a pair of Biolog PM plates of the same type (PM1, PM2A, PM3B, PM4A, or PM9) was taken out of storage (4°C), opened in a biosafety cabinet, and allowed to come to RT. A small disk of dried and UV-irradiated PET (d = 0.458 cm) was added to each of 96 wells in a single PM plate (‘+PET plate’). 100 microL of *P. sakaiensis* in YSV at OD600 = 0.1 were added to each well of the 2 PM plates using either a single-channel repeat or multi-channel pipettor while being careful to not touch the sides or bottom of the wells. The PET disks in the ‘+PET plate’ were plunged to the bottom of each well using a specialty tool made in lab. Each plate was then covered with its lid and sealed with parafilm before being placed in a plate reader or a regular shaker-incubator at 30°C and 260-280 rpm for 4-5 days of growth.

Bacterial growth was monitored via light absorbance (abs600) and recorded every hour (if incubated in a plate reader) or at a single time-point at the end of day 4. A redox indicator (dye G) was then added to each well and plates were incubated for another 24 hrs at 30°C and 260-280 rpm. Metabolic activity was measured at the end of day 5 by photographing plates using a digital camera suspended over a light curtain and calculating the *color depth* in each well (= highest avg. pixel intensity MINUS avg. pixel intensity in well).

Larger PET disks (d = 1.13 cm) were used for downstream screening in 24-well plates. Dilutions of single or combined chemicals/conditions were made in each plate. Plates were closed and sealed with parafilm and left to grow for 2 days in a shaker-incubator at 30°C and 260-280 rpm. Bacterial growth was measured by light absorbance (abs600) at a single time-point at the end of day 2 from 1) total well volumes and 2) 120 microl media samples taken from each well of the 24-well plate(s). The latter report more specifically on planktonic growth of *P. sakaiensis*, while the former also include the absorbance/scattering effects of the PET-bound biofilm and biodegradation-induced changes to PET surface texture and opacity.

### Calculating synergy: growth and metabolic activity scores

Growth and metabolic scores were calculated to discover conditions supporting synergy or enhancement of PET-dependent growth and metabolism using the following formula: (synergy score = delta_readout+PET+x / (delta_readout+PET + delta_readout+x)).

Single time-point measurements were used for calculating both growth and metabolic scores. Multi-time-point growth data were also available for some experiments – in those cases, we used a time-normalized growth integral to calculate growth scores.

The calculation for any given chemical condition requires comparing the relevant observable (for growth or metabolism) from 4 different wells: 1) control with minimal media (YSV) only (negative control); 2) control with YSV media and a single disk of amorphous PET (positive control); 3) well with YSV media and chemical *x*; and 4) well with YSV media, chemical *x*, and a single PET disk. The negative control value is subtracted from 4 to produce the numerator…

The output of the formula reflects whether the combination of chemical *x* and PET had a non-linear effect on bacterial growth or metabolism, with scores above 1 indicating synergistic or greater than expected effects from the chemical*x*:PET combination.

### PET biodegradation assays

Amorphous PET plastic strips were retrieved from 70% ethanol within a biosafety cabinet where they were UV irradiated (10 minutes) and allowed to dry. A pair of PET strips, where individual strips differed slightly in size (each approx. 1.25”x0.31”x0.015”) and shape (round vs. straight edges), was prepared for each condition to be tested. Each pair was submerged in 4 ml of supplemented or unsupplemented (= positive and negative controls) YSV medium in a 15 ml conical and 20 microL of *P. sakaiensis* in YSV at OD600 = 10.05 was added for a final OD600 of 0.05. Cultures were incubated at 30°C with shaking (approx. 270-280 rpm). Every week starting 1 week after the start of the experiment and ending after 5 weeks, a different PET strip (round or straight edges) was taken from each culture and processed (30 minute wash with shaking in 1% SDS, 30 minute wash with shaking in 70% ethanol) and dried before being weighed using an electronic balance (max = 60 g; d = 0.1 mg). Weights were rounded to the nearest mg and recorded and used to calculate biodegradation rates.

### Whole genome sequencing

DNA from a 24 hr culture of *P. sakaiensis* in gm802 was purified using a bacterial DNA kit (Omega Bio-Tek) and prepared into a library for Illumina sequencing by the Alkek Center for Metagenomics and Microbiome Research (CMMR) and the Human Genome Sequencing Center (HGSC) at Baylor College of Medicine.

Sequences were assembled in Geneious Prime (Geneious Prime 2024.0.4) and annotated using RASTtk. Reads were trimmed to a quality score of Q20 and reads that were shorter than 30 bps were filtered out using BBDuk (version 38.84) from the BBMap suite.

## Data availability

Data from PET-dependent bioactivity screening and PET biodegradation assays are provided in the MS in the form of main figures and supplemental figures and tables. The *Piscinibacter sakaiensis* whole genome sequence has been deposited into NCBI with accession number BLANK.

## Acknowledgements

Thanks to NBRC, Japan for providing the lyophilized *Piscinibacter sakaiensis* needed to start this work. Thanks also to Ilse Paola Piedra Aguilar, the wife of FAP, for accidentally initiating this work by showing FAP a digest of the 2016 article by Yoshida et al. introducing *P. sakaiensis* to the world.

## Figures

**Supp Fig 1.**
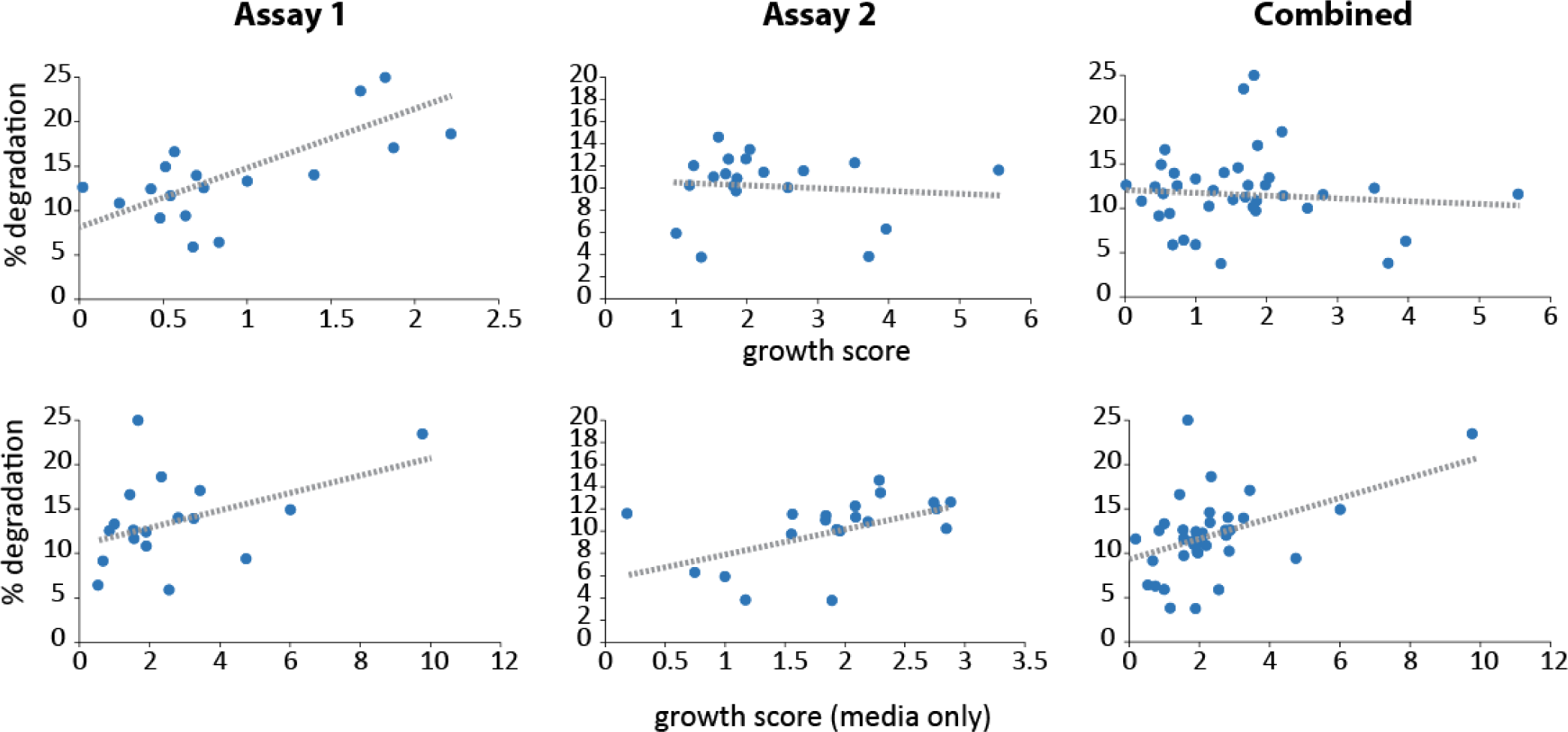
The extent of PET biodegradation after 5 weeks correlates with growth scores. Percent biodegradation after 5 weeks vs. growth scores measured after 2 days for biodegradation assay 1 (left plots), assay 2 (middle plots), and combined data (right plots). Two different growth scores were calculated: 1) from A600 measurements of each well of the 24-well plates (= growth score); and 2) from A600 measurements of 120 microL media taken from each well of the 24-well plates and transferred to a 96-well plate (= media only growth score). A600 measurements from media only are specific for planktonic bacterial growth, while A600 measurements from 24-well plates are affected by variable light scattering from partly degraded PET, the thickness of the PET-bound biofilm, and the presence of planktonic bacteria. (Linear fits, clockwise starting from the top left: 1) y = 5.48x + 8.80; R^2^ = 0.45; 2) y = -0.23x + 10.77; R^2^ = 0.01; 3) y = 0.30x + 12.42; R^2^ = 0.01; 4) y = 1.28x + 9.04; R^2^ = 0.23; 5) y = 2.08x + 6.32; R^2^ = 0.24; and 6) y = 0.98x + 11.18; R^2^ = 0.19).

**Supp Table 1.** Summary of PET biodegradation assay 1. A PET biodegradation assay (see Methods) was performed using 17 test conditions and 2 unsupplemented YSV control conditions [(pos) = with *P. sakaiensis*; (neg) = no bacteria]. Chemical concentrations in each supplemented test condition are given as percentages (w/v) for all but sodium phosphate pH 7 and sodium nitrite (mM). PET strip masses were measured weekly for 5 weeks. The percent biodegradation is the total PET mass degraded by *P. sakaiensis* after 5 weeks. Fold-differences in % biodegradation (fold-diff) were calculated relative to the unsupplemented positive control. An average biodegradation rate (and standard deviation) was calculated from the weekly biodegradation rates for each condition. Fold-differences (fold-diff) were calculated relative to the unsupplemented positive control. Finally, weekly changes in biodegradation rate were calculated and summed to convey the PET biodegradation dynamics supported by each condition.

| condition # | chem1 | [chem1] | % biodegradation | fold-diff | <biodegradation rate> | stdev | fold-diff | weekly change in biodegradation rate (%) |  |  |  |  |
| --- | --- | --- | --- | --- | --- | --- | --- | --- | --- | --- | --- | --- |
|  |  |  |  |  |  |  |  | 1-2 | 2-3 | 3-4 | 4-5 | sum |
| 1 | m-Inositol | 0.13 | 12.43 | 0.93 | 0.090 | 0.020 | 0.94 | 33.14 | -12.37 | -18.49 | -24.50 | -22.23 |
| 2 | m-Inositol | 0.03 | 9.41 | 0.71 | 0.068 | 0.015 | 0.71 | -23.92 | -12.37 | -14.41 | 57.28 | 6.58 |
| 3 | D-Mannitol | 0.13 | 5.89 | 0.44 | 0.042 | 0.040 | 0.44 | -23.92 | -12.37 | -100.00 | 0.00 | -136.29 |
| 4 | L-Serine | 0.50 | 12.62 | 0.95 | 0.091 | 0.009 | 0.94 | 14.12 | -12.37 | 14.12 | -21.36 | -5.50 |
| 5 | L-Serine | 0.02 | 14.03 | 1.05 | 0.102 | 0.043 | 1.06 | -0.15 | 25.19 | -31.53 | -68.54 | -75.04 |
| 6 | GABA | 0.25 | 9.15 | 0.69 | 0.067 | 0.048 | 0.70 | -28.68 | 22.68 | -83.70 | -5.63 | -95.32 |
| 7 | GABA | 0.02 | 11.67 | 0.88 | 0.084 | 0.076 | 0.88 | 128.23 | -48.88 | -67.40 | -100.00 | -88.04 |
| 8 | sodium phosphate | 5.00 | 16.63 | 1.25 | 0.120 | 0.032 | 1.25 | -14.41 | 75.26 | -33.43 | -19.11 | 8.30 |
| 9 | sodium phosphate | 0.63 | 14.93 | 1.12 | 0.108 | 0.014 | 1.13 | -0.15 | 0.15 | -28.68 | 32.12 | 3.44 |
| 10 | sodium nitrite | 5.00 | 6.41 | 0.48 | 0.046 | 0.017 | 0.48 | -14.41 | 16.84 | -71.47 | 183.11 | 114.07 |
| 11 | sodium nitrite | 0.63 | 13.96 | 1.05 | 0.100 | 0.013 | 1.04 | 14.12 | -12.37 | 33.14 | -5.63 | 29.25 |
| 12 | ethylene glycol | 0.25 | 10.83 | 0.81 | 0.077 | 0.067 | 0.81 | 156.76 | -2.63 | -77.18 | -100.00 | -23.05 |
| 13 | ethylene glycol | 0.02 | 12.56 | 0.94 | 0.090 | 0.010 | 0.94 | 14.12 | 2.23 | -18.49 | -5.63 | -7.77 |
| 14 | gm802 | 12.50 | 25.00 | 1.88 | 0.179 | 0.084 | 1.87 | -25.82 | -25.85 | 3.74 | -62.25 | -110.19 |
| 15 | gm802 | 3.13 | 18.64 | 1.40 | 0.135 | 0.073 | 1.41 | -10.34 | -4.40 | -71.47 | 25.83 | -60.38 |
| 16 | gm802 | 0.78 | 17.09 | 1.28 | 0.122 | 0.025 | 1.27 | 33.14 | 0.15 | 14.12 | 17.96 | 65.36 |
| 17 | gm802 | 0.39 | 23.48 | 1.76 | 0.168 | 0.022 | 1.76 | 2.70 | 16.84 | 14.12 | -13.49 | 20.17 |
| 18 (pos) | -- |  | 13.31 | 1.00 | 0.096 | 0.037 | 1.00 | 33.14 | 25.19 | -42.94 | -43.38 | -28.00 |
| 19 (neg) | -- |  | 2.61 | 0.20 | 0.019 | 0.020 | 0.20 |  |  |  |  |  |

**Supp Table 2.**
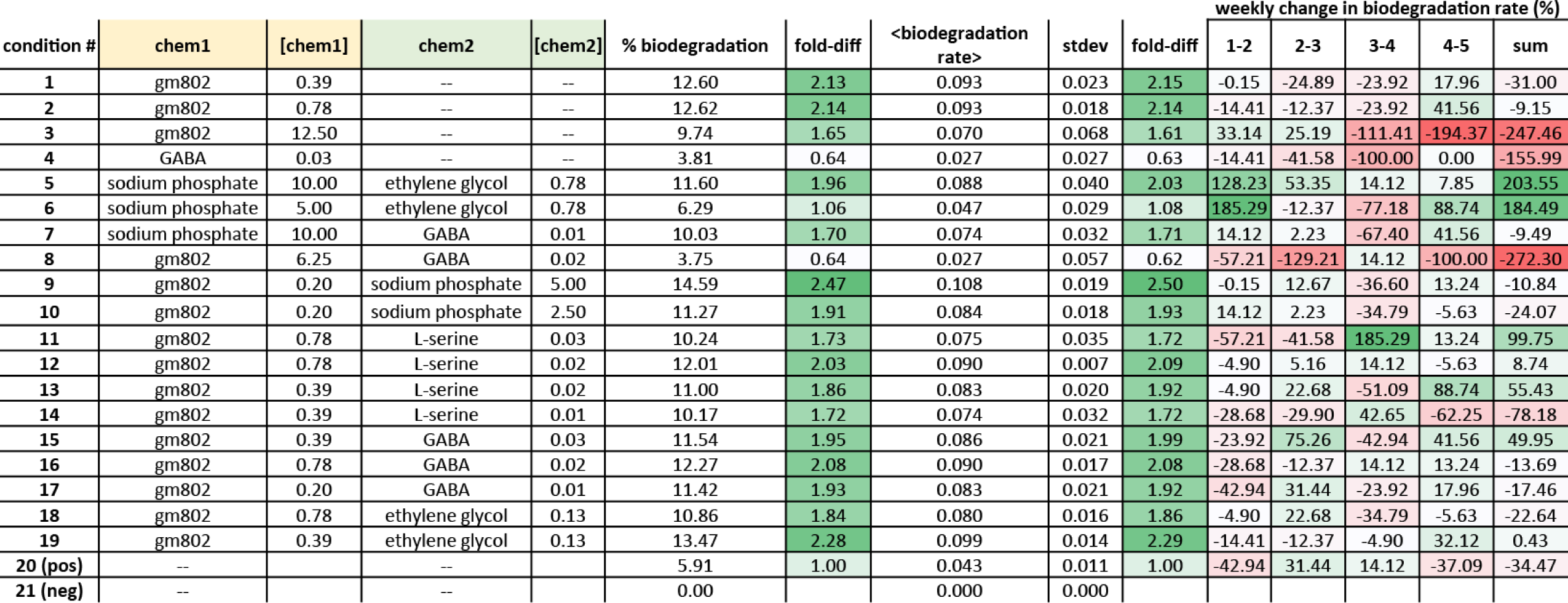
Summary of PET biodegradation assay 2. A PET biodegradation assay (see Methods) was performed using 19 test conditions and 2 unsupplemented YSV control conditions [(pos) = with *P. sakaiensis*; (neg) = no bacteria]. Chemical concentrations in each supplemented test condition are given as percentages (w/v) for all but sodium phosphate pH 7 (mM). PET strip masses were measured weekly for 5 weeks. The percent biodegradation is the total PET mass degraded by *P. sakaiensis* after 5 weeks. Fold-differences in % biodegradation (fold-diff) were calculated relative to the unsupplemented positive control. An average biodegradation rate (and standard deviation) was calculated from the weekly biodegradation rates for each condition. Fold-differences (fold-diff) were calculated relative to the unsupplemented positive control. Finally, weekly changes in biodegradation rate were calculated and summed to convey the PET biodegradation dynamics supported by each condition.

| condition # | chem1 | [chem1] | chem2 | [chem2] | % biodegradation | fold-diff | <biodegradation rate> | stdev | fold-diff | weekly change in biodegradation rate (%) |  |  |  |  |
| --- | --- | --- | --- | --- | --- | --- | --- | --- | --- | --- | --- | --- | --- | --- |
|  |  |  |  |  |  |  |  |  |  | 1-2 | 2-3 | 3-4 | 4-5 | sum |
| 1 | gm802 | 0.39 | -- | -- | 12.60 | 2.13 | 0.093 | 0.023 | 2.15 | -0.15 | -24.89 | -23.92 | 17.96 | -31.00 |
| 2 | gm802 | 0.78 | -- | -- | 12.62 | 2.14 | 0.093 | 0.018 | 2.14 | -14.41 | -12.37 | -23.92 | 41.56 | -9.15 |
| 3 | gm802 | 12.50 | -- | -- | 9.74 | 1.65 | 0.070 | 0.068 | 1.61 | 33.14 | 25.19 | -111.41 | -194.37 | -247.46 |
| 4 | GABA | 0.03 | -- | -- | 3.81 | 0.64 | 0.027 | 0.027 | 0.63 | -14.41 | -41.58 | -100.00 | 0.00 | -155.99 |
| 5 | sodium phosphate | 10.00 | ethylene glycol | 0.78 | 11.60 | 1.96 | 0.088 | 0.040 | 2.03 | 128.23 | 53.35 | 14.12 | 7.85 | 203.55 |
| 6 | sodium phosphate | 5.00 | ethylene glycol | 0.78 | 6.29 | 1.06 | 0.047 | 0.029 | 1.08 | 185.29 | -12.37 | -77.18 | 88.74 | 184.49 |
| 7 | sodium phosphate | 10.00 | GABA | 0.01 | 10.03 | 1.70 | 0.074 | 0.032 | 1.71 | 14.12 | 2.23 | -67.40 | 41.56 | -9.49 |
| 8 | gm802 | 6.25 | GABA | 0.02 | 3.75 | 0.64 | 0.027 | 0.057 | 0.62 | -57.21 | -129.21 | 14.12 | -100.00 | -272.30 |
| 9 | gm802 | 0.20 | sodium phosphate | 5.00 | 14.59 | 2.47 | 0.108 | 0.019 | 2.50 | -0.15 | 12.67 | -36.60 | 13.24 | -10.84 |
| 10 | gm802 | 0.20 | sodium phosphate | 2.50 | 11.27 | 1.91 | 0.084 | 0.018 | 1.93 | 14.12 | 2.23 | -34.79 | -5.63 | -24.07 |
| 11 | gm802 | 0.78 | L-serine | 0.03 | 10.24 | 1.73 | 0.075 | 0.035 | 1.72 | -57.21 | -41.58 | 185.29 | 13.24 | 99.75 |
| 12 | gm802 | 0.78 | L-serine | 0.02 | 12.01 | 2.03 | 0.090 | 0.007 | 2.09 | -4.90 | 5.16 | 14.12 | -5.63 | 8.74 |
| 13 | gm802 | 0.39 | L-serine | 0.02 | 11.00 | 1.86 | 0.083 | 0.020 | 1.92 | -4.90 | 22.68 | -51.09 | 88.74 | 55.43 |
| 14 | gm802 | 0.39 | L-serine | 0.01 | 10.17 | 1.72 | 0.074 | 0.032 | 1.72 | -28.68 | -29.90 | 42.65 | -62.25 | -78.18 |
| 15 | gm802 | 0.39 | GABA | 0.03 | 11.54 | 1.95 | 0.086 | 0.021 | 1.99 | -23.92 | 75.26 | -42.94 | 41.56 | 49.95 |
| 16 | gm802 | 0.78 | GABA | 0.02 | 12.27 | 2.08 | 0.090 | 0.017 | 2.08 | -28.68 | -12.37 | 14.12 | 13.24 | -13.69 |
| 17 | gm802 | 0.20 | GABA | 0.01 | 11.42 | 1.93 | 0.083 | 0.021 | 1.92 | -42.94 | 31.44 | -23.92 | 17.96 | -17.46 |
| 18 | gm802 | 0.78 | ethylene glycol | 0.13 | 10.86 | 1.84 | 0.080 | 0.016 | 1.86 | -4.90 | 22.68 | -34.79 | -5.63 | -22.64 |
| 19 | gm802 | 0.39 | ethylene glycol | 0.13 | 13.47 | 2.28 | 0.099 | 0.014 | 2.29 | -14.41 | -12.37 | -4.90 | 32.12 | 0.43 |
| 20 (pos) | -- |  | -- |  | 5.91 | 1.00 | 0.043 | 0.011 | 1.00 | -42.94 | 31.44 | 14.12 | -37.09 | -34.47 |
| 21 (neg) | -- |  | -- |  | 0.00 |  | 0.000 | 0.000 |  |  |  |  |  |  |

**Supp Table 3.**
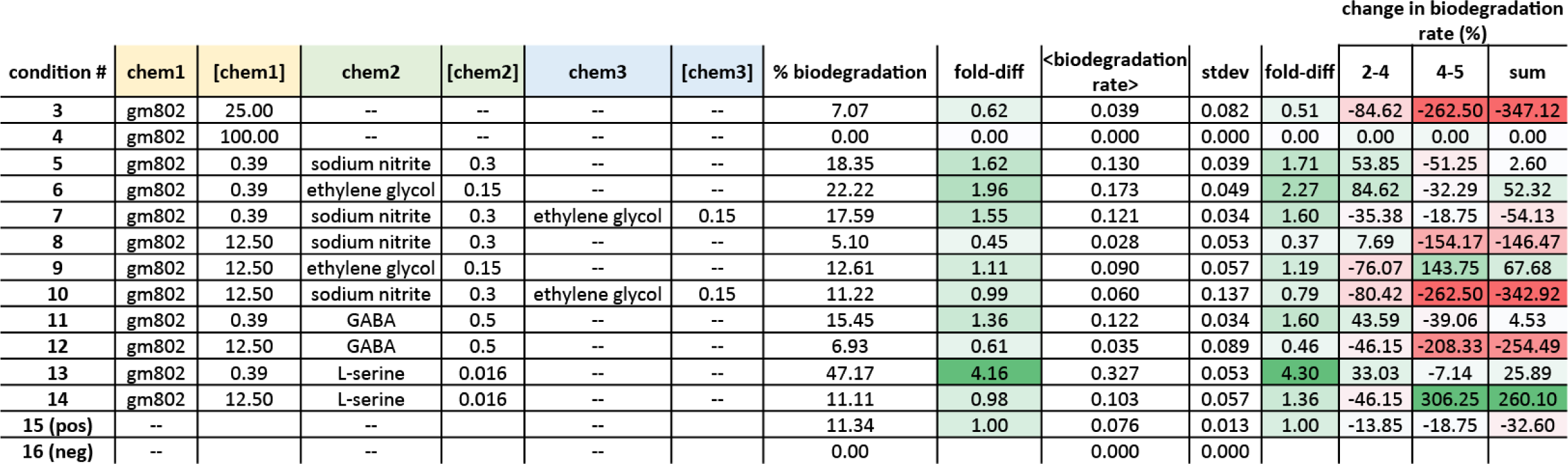
Summary of PET biodegradation assay 3. A PET biodegradation assay (see Methods) was performed using 14 test conditions and 2 unsupplemented YSV control conditions [(pos) = with *P. sakaiensis*; (neg) = no bacteria]. Chemical concentrations in each supplemented test condition are given as percentages (w/v) for all but sodium nitrite (mM). PET strip masses were measured at 2, 4, and 5 weeks. The percent biodegradation is the total PET mass degraded by *P. sakaiensis* after 5 weeks. Fold-differences in % biodegradation (fold-diff) were calculated relative to the unsupplemented positive control. An average biodegradation rate (and standard deviation) was calculated from the 3 biodegradation rates for each condition. Fold-differences (fold-diff) were calculated relative to the unsupplemented positive control. Finally, changes in biodegradation rate between weeks 2 and 4 and 4 and 5 were calculated and summed to convey the PET biodegradation dynamics supported by each condition.

| condition # | chem1 | [chem1] | chem2 | [chem2] | chem3 | [chem3] | % biodegradation | fold-diff | <biodegradation<br>rate> | stdev | fold-diff | 2-4 | 4-5 | sum |
| --- | --- | --- | --- | --- | --- | --- | --- | --- | --- | --- | --- | --- | --- | --- |
| 3 | gm802 | 25.00 | -- | -- | -- | -- | 7.07 | 0.62 | 0.039 | 0.082 | 0.51 | -84.62 | -262.50 | -347.12 |
| 4 | gm802 | 100.00 | -- | -- | -- | -- | 0.00 | 0.00 | 0.000 | 0.000 | 0.00 | 0.00 | 0.00 | 0.00 |
| 5 | gm802 | 0.39 | sodium nitrite | 0.3 | -- | -- | 18.35 | 1.62 | 0.130 | 0.039 | 1.71 | 53.85 | -51.25 | 2.60 |
| 6 | gm802 | 0.39 | ethylene glycol | 0.15 | -- | -- | 22.22 | 1.96 | 0.173 | 0.049 | 2.27 | 84.62 | -32.29 | 52.32 |
| 7 | gm802 | 0.39 | sodium nitrite | 0.3 | ethylene glycol | 0.15 | 17.59 | 1.55 | 0.121 | 0.034 | 1.60 | -35.38 | -18.75 | -54.13 |
| 8 | gm802 | 12.50 | sodium nitrite | 0.3 | -- | -- | 5.10 | 0.45 | 0.028 | 0.053 | 0.37 | 7.69 | -154.17 | -146.47 |
| 9 | gm802 | 12.50 | ethylene glycol | 0.15 | -- | -- | 12.61 | 1.11 | 0.090 | 0.057 | 1.19 | -76.07 | 143.75 | 67.68 |
| 10 | gm802 | 12.50 | sodium nitrite | 0.3 | ethylene glycol | 0.15 | 11.22 | 0.99 | 0.060 | 0.137 | 0.79 | -80.42 | -262.50 | -342.92 |
| 11 | gm802 | 0.39 | GABA | 0.5 | -- | -- | 15.45 | 1.36 | 0.122 | 0.034 | 1.60 | 43.59 | -39.06 | 4.53 |
| 12 | gm802 | 12.50 | GABA | 0.5 | -- | -- | 6.93 | 0.61 | 0.035 | 0.089 | 0.46 | -46.15 | -208.33 | -254.49 |
| 13 | gm802 | 0.39 | L-serine | 0.016 | -- | -- | 47.17 | 4.16 | 0.327 | 0.053 | 4.30 | 33.03 | -7.14 | 25.89 |
| 14 | gm802 | 12.50 | L-serine | 0.016 | -- | -- | 11.11 | 0.98 | 0.103 | 0.057 | 1.36 | -46.15 | 306.25 | 260.10 |
| 15 (pos) | -- |  | -- |  | -- | -- | 11.34 | 1.00 | 0.076 | 0.013 | 1.00 | -13.85 | -18.75 | -32.60 |
| 16 (neg) | -- |  | -- |  | -- | -- | 0.00 |  | 0.000 | 0.000 |  |  |  |  |

